# Clustering single cell CITE-seq data with a canonical correlation based deep learning method

**DOI:** 10.1101/2021.09.07.459236

**Authors:** Musu Yuan, Liang Chen, Minghua Deng

## Abstract

Single cell sequencing examines the sequence information from individual cells with optimized next generation sequencing (NGS) technologies. It provides researchers a higher resolution of cellular differences and a better understanding of the function of an individual cell in the context of its microenvironment. CITE-seq, or Cellular Indexing of Transcriptomes and Epitopes by sequencing, is one of the latest innovations in the domain of single cell sequencing. It enables researchers to simultaneously capture RNA and surface protein expression on the same cells so that we can correlate the two data types, identify biomarkers and better characterize cell phenotypes. Although multi-omics sequencing technologies developed rapidly, data analyzing methods tailored for multiomics sequencing data are lacking. Several serious problems have to be faced. An important one is how to integrate the information from different modalities, namely scRNA and protein data, efficiently.

In this paper, we introduce a canonical correlation based deep learning method called scCTClust for clustering analysis over CITE-seq data. We impute and extract the characteristics of the high dimensional RNA part of data with a ZINB model-based autoencoder. A t-kernel distance is introduced to measure the similarity between query cell and cluster centroids. And the protein data rectifies the feature extraction of scRNA data in a canonical correlation analysis(CCA) way. Extensive data experiments shows that scCTClust can precisely recover the dropout values for RNA sequencing data and extract authentic information from both modalities, getting a much better clustering result than state-of-the-art methods, no matter single-omic or multi-omics clustering algorithms.

## 1. Introduction

High-throughput single-cell sequencing technology represented by single-cell RNA sequencing has been widely used in cancer tumors and embryonic development in recent years. The gene expression profile obtained by transcriptome sequencing allows us to study tissue heterogeneity at the individual cellular level. Although the amount of sequencing data is increasing, single-cell RNA sequencing data still meet the characteristics of high noise and dimensions, which limits the accuracy of the calculation method to a certain extent. The recent rise of single-cell multi-omics sequencing technology has allowed researchers to obtain information on the epigenomics, genetics and proteomics of individual cells at the same time. The expression profiles of these molecules provide us with a more comprehensive and in-depth view of cell states and interactions. Among the various single-cell multi-omics sequencing technologies, cellular indexing of transcriptomes and epitopes by sequencing (CITE-seq) [12] allows simultaneous quantification of gene expression and surface proteins by using single-cell RNA-sequencing (scRNA-seq) and antibody-derived tags (ADTs) in single cells. Compared with gene expression profiles, surface protein expression profiles hold weaker sparsity and lower dimensionality. In this article, we focus on the integrated analysis of transcriptomics and proteomics based on CITE-seq.

The identification of cell types plays an essential role in the analysis of single-cell sequencing data. It is inseparable from discovering key regulatory genes and digging meaningful cell pathways. In the past few years, clustering methods for single-cell RNA sequencing data have emerged in endlessly, including graph segmentation based (Seurat…), class center based (RaceID…) [6], ensemble learning based (SC3…) and deep learning based (scziDesk…) [3]. These customized methods have not been well extended to multiomics cluster analysis due to the limitations of modeling. Naturally, the accuracy of cell type discrimination will also be affected. How to integrate and statistically model the relationship between RNA molecular expression profile and protein molecular expression profile to further improve the effect of single-cell multi-omics clustering is worthy of further exploration and research.

Human cells contain about 20,000 genes, each cell has its own specific gene expression pattern, only some genes are expressed, resulting in cell-specific protein components and biological functions. The recent springing up of single cell RNA sequencing(scRNA-seq) technology has enabled researchers to study gene expression patterns at the single cell level, thus enabling more accurate research on intercellular heterogeneity. Extending scRNA-seq, cellular indexing of transcriptomes and epitopes by sequencing (CITE-seq) simultaneously measures the abundance of RNA and selected proteins on the cell surface with results comparable to gold-standard flow cytometry. The advent of CITE-seq provides us an exciting opportunity to utilize multi-omics data to characterize the specificity of single cells more stereoscopically in many ways.

In CITE-seq experiment, the abundance of RNA and surface marker is quantified by Unique Molecular Index (UMI) [11] and Antibody-Derived Tags (ADT) respectively, for a common set of cells at the single cell resolution. These two data sources represent different but highly related and complementary biological components. Though both data were widely used in cell type identification, current statistical methods for jointly analyzing data from scRNA-Seq and CITE-Seq are still unavailable or immature and most of which are conventional ones, such as MOFA [2], BREMSC [19], jointDIMMSC [14], offering no solution for big data cases.

For jointly clustering of multi-omics data from CITE-seq, we have to resolve several problems facing. During the reverse transcription and the amplification step, RNA transcripts might be missed and consequently not detected in the following sequencing. This is called the dropout problem. One mature deep learning method for clustering over scRNA-seq data is called scDeepCluster [16] which integrates the ZINB(zero-inflated negative binomial) model with clustering loss in a principled way. Roughly speaking, a low-dimensional latent representation of RNA-seq data is learned by the ZINB model-based autoencoder, while the clustering task on latent space is performed by clustering with Kullback–Leibler (KL) divergence, as described in the ‘deep embedded clustering’ (DEC) [20] algorithm.

Another challenge we are facing is how to integrate informations of different modalities. Deep Canonical Correlation Analysis(DCCA) [1] is a method to learn complex nonlinear transformations of two views of data such that the resulting representations are highly linearly correlated. However, methods alike has never been introduced in to the fields of bioinfomatics.

Here, we introduced a CCA-based deep learning clustering method for CITE-seq data called scCTClust. We apply a denoising stacked autoencoder [17] and ZINB model to RNA count matrix to eliminate the effects of dropout events along with dimension reduction. In the proposed framework, the nonlinear function mapping the RNA count matrix to a low-dimensional latent representation is learned by the ZINB(zero-inflated negative binomial) model-based autoencoder. We introduce and optimize a CCA loss to maximize the correlation between the latent representation of RNA data and protein counts data. Then we cluster on the latent layer of the autoencoder. In that way, the protein part of data provides auxiliary information to correct characteristics extraction of RNA data and improves clustering results. Extensive real data and simulation data experiments prove that, scCTClust surpassed all competing methods comprehensively.

## 2. Results

In this section, we first introduce the real and simulated datasets we used, then we show the hyperparameters setting of scCTClust model. We compare scCTClust with other state-of-the-art methods and conduct series of self-contrast experiments to illustrate the effectiveness and efficiency of scCTClust. We also design series of simulation experiments to explore whether our methods is highly adaptable. In cluster analysis, three index are often introduced to evaluate the effectiveness of a method, adjusted mutual information (AMI) [13], clustering accuracy (CA) and adjusted Rand index (ARI) [9]. The ranges of AMI and CA are from 0 to 1, while the ARI can yield negative values. The three metrics are statistics of the concordance of two clustering labels; the higher the values, the higher concordance of the clustering. As random seed are necessity in the stage of simulation data generating and network training, we repeat the experiments for 10 times under different seeds to make sure that our results is convincing and not out of incident.

### 2.1. Parameter setting

The hyperparameters of scCTClust model contains learning rate, batch size, standard deviation of noise, training epoches(pretrain, midtrain, funetrain), scale parameters (*α, γ*) adjusting the composition of loss function, as well as neural numbers (dimensions) in encoder layers. In all real data experiments, we fixed the learning rate = 1*e* − 4, batch size = 2000, noise_sd = 1.5, training epoches = (600, 100, 200) and dimensions of encoder layer = [256, 64, 32] across different datasets. The only hyperparameter tuned below is the scale factor (*α, γ*).

### 2.2. datasets

As Cite-seq becomes widely used in cases including finding differential expression genes and clustering analysis, many labs cast their sights on utilizing CITE-seq data for a more comprehensive insights into the biological systems under study, thus produced a great deal of single cell CITE-seq data labeled under gold-standard. The real datasets we are going to operate our method on afterwards are mostly produced in this way.

We use a published cord blood mononuclear cells (CBMCs) dataset used by Seurat for multimodal vignettes and can be downloaded from Satija Lab website (https://satijalab.org/seurat/v3.2/multimodal_vignette.html). A total of 7,895 cells containing 20501 different genes from a healthy donor were stained with 10 markers and divided into 8 different cell types under golden standard. We also applied all these methods to a group of murine spleen and lymph node cells, which contain 13553 heterogeneous immune cell populations that are well-characterized by 110 surface protein markers. This group of data was produced by Gayoso’s group and was used in their article *A Joint Model of RNA Expression and Surface Protein Abundance in Single Cells* [5]. Its CITE-seq experiments were executed on the 10x Genomics Chromium microfluidic platform using BioLegend TotalSeq-A antibodies.

### 2.3. Comparison with other methods

We compare our scCTClust with three state-of-the-art CITE-seq analyzing methods, Multi-Omics Factor Analysis(MOFA), Bayesian Random Effects Mixture Model(BREMSC) and Dirichlet mixture model for clustering droplet-based single cell transcriptomic data(JointDIMMSC). We also run a scRNA-seq data based method scDeepCluster [16] on scRNA-seq part of the CITE-seq data used in following experiments.

We apply those five methods over CBMC, CBMC(f), SLN, SLN(f), where “f” represents filtered dataset in which 500 highly variable genes are selected. First of all, over all datasets, there are essential differences between the running time of scCTClust and conventional methods MOFA, BREMSC and jointDIMMSC. Namely, with an Nvidia Geforce RTX TITAN(24G), the average running time of scCTClust with ten different random seeds is 440.89s, while for those three conventional methods running time are all over 3 hours. Under the evaluation of CA, AMI and ARI, scCTClust outperforms all other state-of-the-art methods over two datasets, showing its dominant position in the field of CITE-seq cluster analysis, especially for big data cases. And we believe that with out selecting highly variable genes which reduce the dimension of input data, the gap between scCTClust and competing methods would surely expand. Full results are shown in details in Figure.1 in form of bar plots.

**Figure 1.**
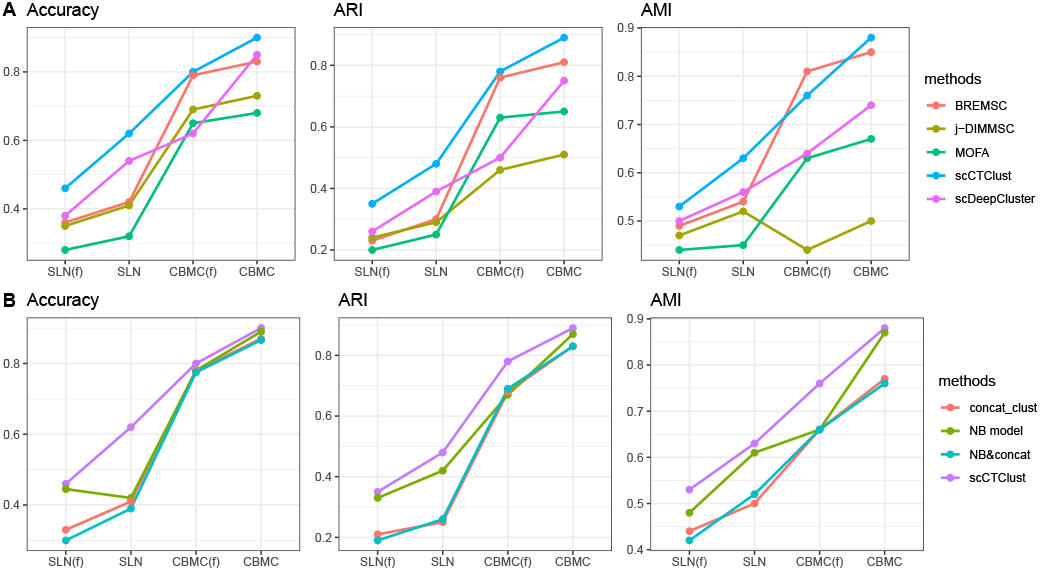
Realdata experiment results. Figure A is the line chart of five algorithms’ performance over SLN(f), SLN, CBMC(f) and CBMC datasets, evaluated by accuracy, ARI and AMI from left to right. Figure B is the line chart of self-contrast experiments results over the same four results.

**Figure 2.**
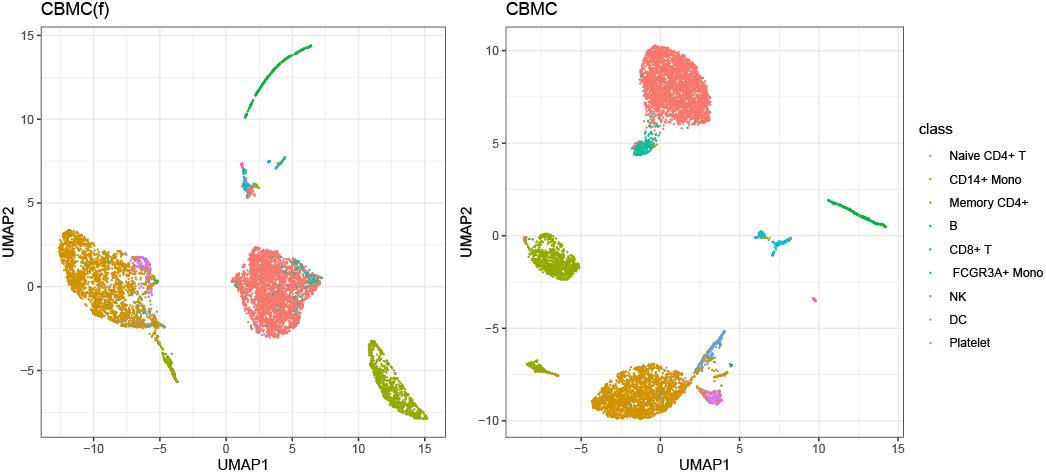
Real data experiment UMAP. Two-dimensional UMAP visualization plots of cell types for scCTClust over CBMC(f) and CBMC.

**Figure 3.**
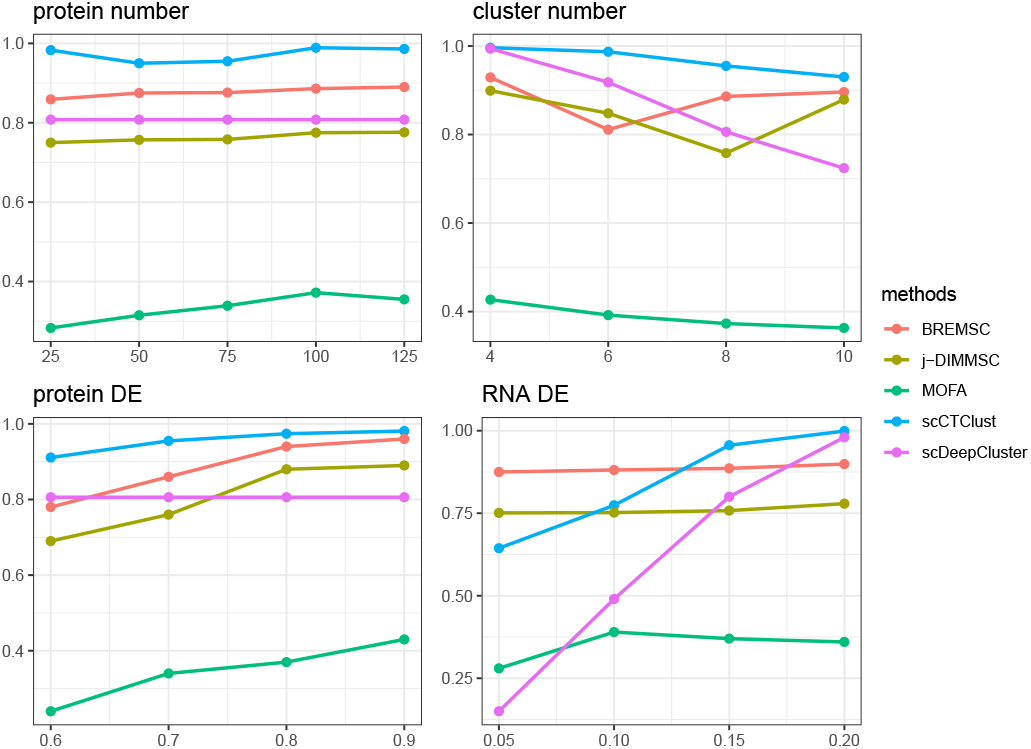
Simulation experiment results. Line charts for five methods in simulation experiments. X label represents the value of subfigure’s title. Y label represents the value of accuracy.

### 2.4. Some self-contrast experiments

In this section, we design series of self-contrast experiments for scCTClust over CBMC, CBMC(f), SLN and SLN(f). We apply five methods on all four datasets. scCTClust surpasses all competing methods under evaluation index including CA, ARI and AMI. Detailed results of self-contrast experiments are shown in Fig.1. First, we explore which is better, to cluster over the concatenated layer or merely use protein data as an auxiliary optimizing the construction of RNA data representation through cca loss and operate clustering on optimized RNA representation. According to our experiments, scCTClust takes a heavy lead over all datasets, indicating that using protein data as auxiliary message works better than executing clustering over the concatenate layer of RNA and protein representation matrix.

Second, we consider if there is a meaningful distinct between ZINB and NB model at dropout events alignment. We replace the ZINB loss with NB loss in scCTClust and maintain all other parts of their model. We also replace the ZINB loss with NB loss for the derivative model which clusters over the concatenated layer as an extra experiment. According to a recent article *Droplet scRNA-seq is not zero-inflated* [15], the widely utilize of ZINB model while handling droplet based RNA-seq data is totally unnecessary. Our results is that scCTClust with ZINB model gained an advantage that can’t be ignored comparing to the one with NB model over CBMC and Spleen&Lymph cell datasets. Moreover, ZINB model gains great advantages in simulation experiments conducted in case studies, thus we hold the view that ZINB model plays an important part in RNA data characterizing and feature extracting.

Third, for clustering stage, we explore whether the ‘cdec’ loss we introduced above evidently reduce the clustering time and boost clustering performance. We compare the CA, ARI, AMI and convergence time among scCTClust adopting 5 different clustering losses including “square”, “L21”, “sphere” [21], “dec” and “cdec” loss. The first three separately refers to soft kmeans loss based on Euclid distance, L21 norm distance and sphere distance. The comparison measured by AC, AMI, ARI and convergence time is shown in details in Table.1. The results indicate that while cdec occupies the fist rank in the comparison by CA, AMI and ARI, its convergence time is only slightly longer than square loss, far less than L21, sphere and dec loss. With no doubt the results strongly prove the superiority of cdec loss in the case of clustering over CITE-seq data latent representation learned by our model.

**TABLE 1.** Performance of scCTClust with different clustering loss.

|  | square | L21 | sphere | dec | cdec |
| --- | --- | --- | --- | --- | --- |
| CA | 0.8765 | 0.8627 | 0.7811 | 0.8728 | 0.9074 |
| ARI | 0.8365 | 0.8497 | 0.7493 | 0.8498 | 0.8852 |
| AMI | 0.7535 | 0.7622 | 0.6842 | 0.7718 | 0.7834 |
| Time | 3.856 | 6.526 | 24.03 | 16.08 | 4.154 |

**TABLE 2.** Simulation data settings For all experiments, each cluster contains 500 cells and we only varies one of other four parameters, marked as ‘*’ in the table.

| Cell/Cluster | Cluster | Protein | Prob<sub>RNA</sub> | Prob<sub>protein</sub> |
| --- | --- | --- | --- | --- |
| 500 | * | 75 | 0.15 | 0.7 |
| 500 | 8 | * | 0.15 | 0.7 |
| 500 | 8 | 75 | * | 0.7 |
| 500 | 8 | 75 | 0.15 | * |

As the latent space in the proposed scCTClust model is an ideal low-dimensional embedded representation of the high-dimensional input data. To illustrate the representation effectiveness of latent space, we apply UMAP to visualize the feature embeddings of scCTClust, by mapping them to a two-dimensional space. In CBMC and CBMC(f) experiments, scCTClust separates different cell types clearly, especially for big cell types with abundant query samples such as “Naive CD4+ T”, “CD14+ Mono” and “Memory CD4+”. The UMAP plots gives us a more intuitive look of scCTClust’s powerful clustering analyzing ability for CITE-seq data.

### 2.5. Case studies

In this section, we generate following datasets to simulate CITE-seq data acquired from cells in different biological scenarios. All simulation data was generated utilizing R package splatter.

Specifically, We aim to investigate the performance of scCTClust against competing methods under different cluster numbers, protein data dimensions and differential expression features probabilities. We fix the number of every group of cells at 500 during our experiments. We investigate the performance of each method under different cluster numbers which varied in 4,6,8,10 while gene numbers and protein types fixed at 2500 and 75 differential expression genes/proteins probabilities at (0.15, 0.7). Moreover, under the guidance of control variate technique, we conduct series of simulation experiments, the details of which are listed in Table.**??**.

We first investigate the performance of each method under different cluster numbers which varies in 4,6,8,10. Under all circumstances, scCTClust keeps an obvious lead in CA, AMI and ARI, fully manifests the superiority of scCTClust handling CITE-seq data troubled by dropout events and multi-omics information integration. Moreover, comparing the decrease scale of CA when shifting the cluster number from 4 to 10. The decrease of CA for scCTClust is 6.67% while BREMSC, jointDIMMSC and scDeepCluster at least take a 10% decline in accuracy. It shows the great adaptability of scCTClust when processing big sequencing data.

We next evaluate the performance of according methods with different numbers of protein types. We vary the number of different types of surface protein from 25 to 125 to investigate whether the number of protein types matters to the performance of scCTClust. The reason why we select the interval [25, 125] to execute the experiments over was that under most cases we are incapable of sequencing more than several hundreds of types of proteins simultaneously at single cell level. For all methods, running time increased noticeably as the number of different protein types increasing but AC, ARI and AMI did not. Under all circumstances, scCTClust surpass all other state-of-the-art multi-omics clustering methods, including BREMSC, jointDIMMSC and MOFA, significantly providing a strong evidence that scCTClust have a better performance on CITE-seq clustering analysis no matter how many types of surface protein contained. However, as the dimension of protein data grows, the performance of scCTClust does not get better linearly. A plausible explanation may be the curse of dimension faced during the clustering process.

Finally, we investigate the performance of scCTClust while varying differential expression features probabilities. Differential expression features probabilities refers to differential expression genes probabilities and differential expression proteins probabilities, by adjusting the de_prob parameters in splatter, it is possible for us to control the simulating RNA and protein data’s quality so that we can make an inquiry if scCTClust is capable of adapting all qualities of datasets. The detailed evaluation of clustering performance was measured by AC, ARI and AMI. On the first half, we fix the RNA de_prob 0.15 and change the protein de_prob from 0.6 to 0.9 to investigate the changes in relating methods’ performance. For all methods, performance improves ordinarily while the de_prob growing. And the performance of scCTClust is beyond all others as we expected, undoubtedly demonstrating its powerful effect over multi-modal clustering analysis.

On the other half, we vary the de_prob for RNA analogue data by 0.05/0.1/0.15/0.2 and fix the de_prob for protein data 0.7 to simulate the case that the quality of RNA data fluctuates while protein data quality is high. When RNA de_prob is over 0.1, scCTClust and its derivative methods took a major lead on clustering performance, indicating that when RNA data is of high quality, scCTClust can effectively utilize protein data to ameliorate clustering process. While RNA de_prob is severely low, the power of scCTClust is weaker than conventional methods. In considering of the even worse performance scDeepCluster gained under same situations, a probable reason may be that the ZINB-based autoencoder is not capable of extract the character of data effectively when RNA data is under such poor conditions. The performance of scCTClust increases along with the increasing of de_prob for RNA part of simulating data which behaved as we expected. For all simulation experiments, detailed AC, ARI, AMI results are displayed in Fig.1.

## 3. Methods

### 3.1. Read count data pre-processing

A *n* cells raw CITE-seq read count data which consists of a RNA count matrix with *p* genes and a ADT count matrix with *r* proteins are processed by the Python package SCANPY. First, features with no count in any cell are filtered out. Second, size factors are calculated and read counts are normalized by library size, so total counts are same across cells in each matrix. Formally, if we denote the features library size (number of total read counts) of cell *i* as *s*_*i*_, then the size factor of cell *i* is *s*_*i*_/median(*s*). The last step is to take the log transform and scale of the read counts, so that count values in each matrix follow unit variance and zero mean. The pre-processed RNA count matrix is treated as the input for our denoising ZINB model-based autoencoder while the pre-processed ADT count matrix to be concatenated with the latent layer of the autoencoder. The architecture of model scCTClust is shown in Fig.4.

**Figure 4.**
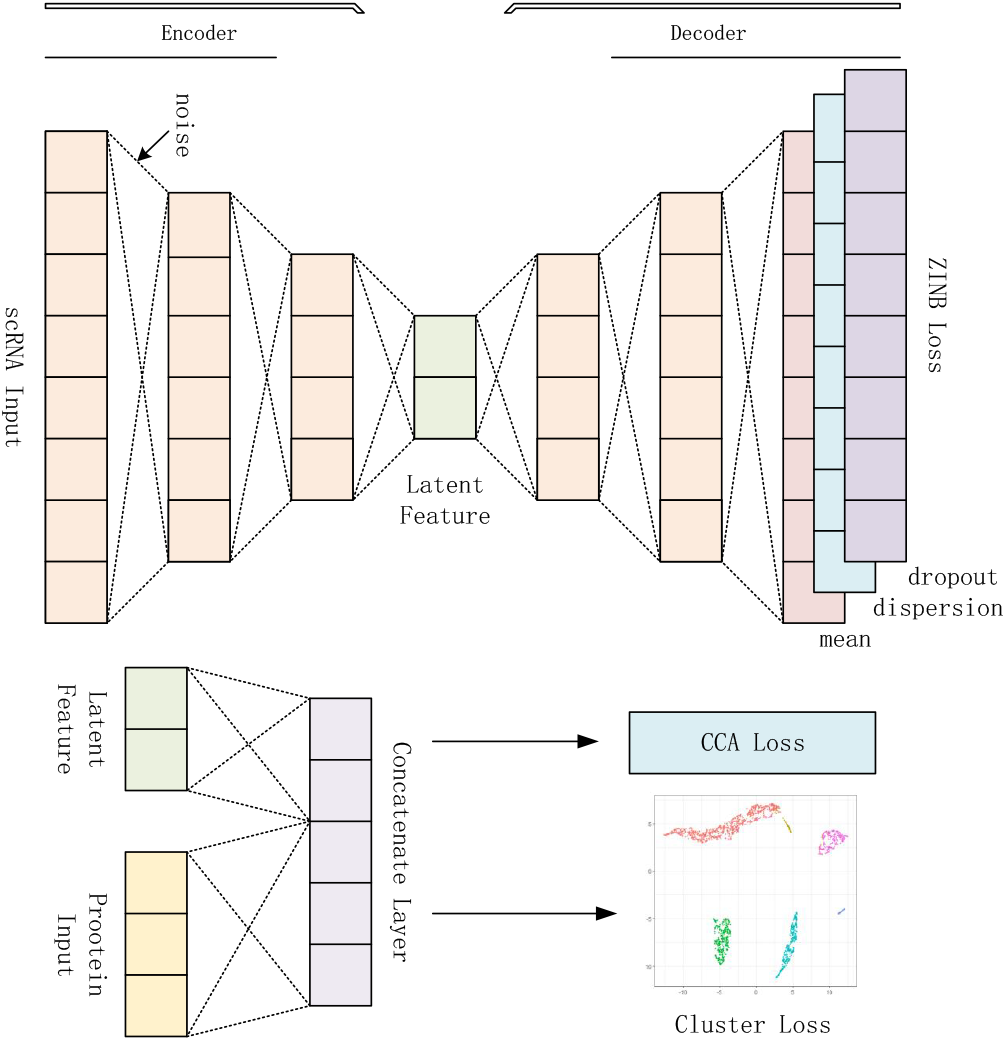
Framework Overview.

### 3.2. Denoising ZINB model-based autoencoder

The autoencoder is a type of artificial neural network used to extract efficient feature representation in an unsupervised manner. The denoising autoencoder is an autoencoder that receives corrupted data points as input and is trained to predict the original incorrupt data points as its output. The autoencoder usually has a low-dimensional bottleneck layer to learn a latent feature representation, referring to cells’ genes character in this case. The denoising autoencoder is proved to be much more powerful in learning a robust representation of data, because it has the ability to learn the representations of the input that are corrupted by small irrelevant changes in the input. Here, we apply the denoising autoencoder technique to map the input of RNA read counts to an embedded space to carry out concatenating and clustering better. In practice, we first corrupt the RNA input with random Gaussian noise, then construct the autoencoder with regular fully connected layers. Formally, RNA input *X* is corrupted by noise

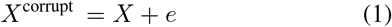

where *e* represents the random Gaussian noise. Note that the noise can be incorporated into every layer of the encoder, which is defined as a stacked denoising autoencoder. We define the encoder function as *z* = *f*_*w*_ (*X*^corrupt^) and the decoder function *X′* = *g*_*W′*_ (*z*). The encoder and decoder functions are both fully connected neural networks with rectifier activation. Here *W* and *W′* are the learned weights of the functions. The learning process of the denoising autoencoder minimizes the loss function

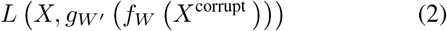

where *L* is the loss function.

To capture the character of RNA-seq data, we apply a ZINB(zero-inflated negative binomial) model-based autoencoder instead of regular stacked autoencoder, which is trained to attempt to reconstruct its input. Unlike the regular autoencoder, the loss function of the ZINB model-based autoencoder is the likelihood of a ZINB distribution. ZINB is applied to characterize the dropout events in scRNA-seq [4]. At this case, a wide application of NB(negative binomial) model when handling scRNA-seq data is worth mentioning. But we still apply ZINB model to RNA data in our method as it fits the data better and gets great clustering results which will be shown by details in latter section. Formally, ZINB is parameterized with the mean (*µ*), the dispersion (*θ*) of the negative binomial distribution and with an additional coefficient (*x*) that represents the weight of the point mass of probability at zero (the probability of dropout events):

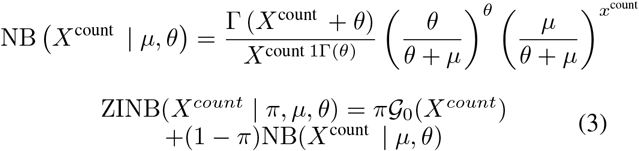

where *X*^count^ represents the raw read counts. The ZINB model-based autoencoder estimates the parameters *µ, θ* and *π*. If 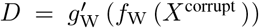 represents the last hidden layer of decoder, we append three independent fully connected layers to *D*. to estimate the parameters

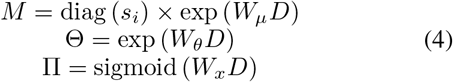

where *M*, Θ and Π represent the matrix form of estimations of mean, dispersion and dropout probability, respectively. The size factors *s*_*i*_ are calculated in the ‘data pre-process’ part and are included as an independent input to the deep learning model. The activation function chosen for mean and dispersion is exponential because the mean and dispersion parameters are non-negative values. The loss function of the ZINB model-based autoencoder is the negative log of the ZINB likelihood:

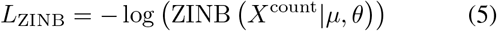

### 3.3. Canonical concatenate analysis

For typical canonical concatenate analysis without deep neural networks [8], Let 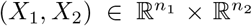 denote random vectors with corvariances (Σ_11_, Σ_22_) and crosscorvariance Σ_12_. CCA finds pairs of linear projections of the two views 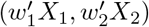 that are maximally correlated:

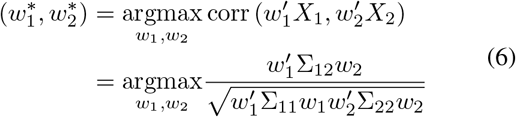

since the objective is invariant to scaling of *w*_1_ and *w*_2_ the projections are constrained to have unit variance:

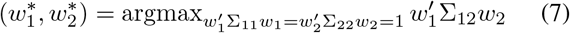

When finding multiple pairs of vectors 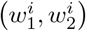, subsequent projections are also constrained to be uncorrelated with previous ones, that is 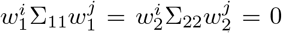 for *i < j*. Assembling the top *k* projection vectors 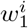 into the columns of a matrix 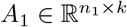, and similarly placing 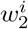 into 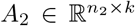 we obtain the following formulation to identify the top *k* ≤ min (*n*_1_, *n*_2_) projections:

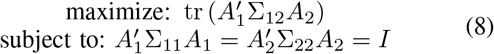

There are several ways to express the solution to this objective; we follow the one in [10]. Define 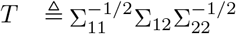, and let *U*_*k*_ and *V*_*k*_ be the matrices of the first *k* left- and right-singular vectors of *T*. Then the optimal objective value is the sum of the top *k* singular values of *T* (the Ky Fan *k* -norm of *T*) and the optimum is attained at

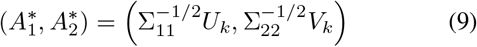

Note that this solution assumes that the covariance matrices Σ_11_ and Σ_22_ are nonsingular, which is satisfied in practice because they are estimated from data with regularization: given centered data matrices 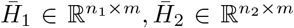, one can estimate, e.g.

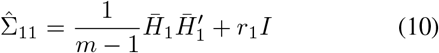

where *r*_1_ *>* 0 is a regularization parameter.

Back to our scCTClust model, from the latent layer of the denoising ZINB-model based autoencoder, we get a low-dimension representation of the RNA read count matrix which contains all information of RNA sequencing data. Let 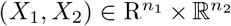 denote the RNA input and protein input of the autoencoder, *H*_1_ = *f* (*X*_1_) denote the output of autoencoder’s latent layer where *f* a non-linear mapping defined by the encoder part of the autoencoder. In most cases, the dimension of protein data matrix is quite low, often a hundred or less, so we consider dimension reduction for protein matrix as unnecessary and concatenate the protein data with the latent representation of RNA data instantly, which means *H*_2_ = *X*_2_. At this stage, we aim to add a CCA_loss to the ZINB loss function of the denoising autoencoder,

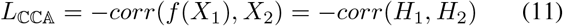

to find an adequate representation of RNA read count matrix which is maximally correlated with the ADT read count matrix while maintaining the autoencoder’s denoising performance for RNA data.

To find a simple presentation of *L*_*CCA*_, we assume our dataset contains *n* cells and the dimension of autoencoder’s latent layer is *o*. Then *H*_1_ ∈ ℝ^*o*×*n*^, *H*_2_ ∈ ℝ^*r*×*n*^ indicate the representation of RNA count matrix and protein count matrix separately. Let 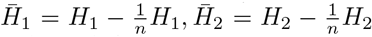 be the centered data matrix, and define

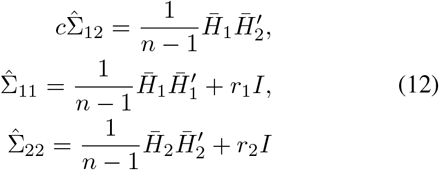

where *r*_1_, *r*_2_ are regularization constants. Assume that *r*_1_ *>* 0, *r*_2_ *>* 0 so that 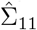 and 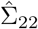 are positive definite.

As is discussed ahead for CCA, the total correlation of the top *k* components of *H*_1_ and *H*_2_ is the sum of the top *k* singular values of the matrix 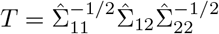 If we use all components then this is exactly the matrix trace norm of *T*, or

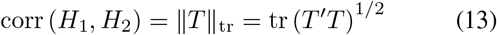

As our input data is CITE-seq data which means the sizes of 2 data matrices are different under most circumstances, we calculate the correlation by compute the top *k* singular values of the matrix 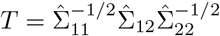 utilizing singular value decomposition(SVD) [18].

### 3.4. Deep embedded clustering with local structure preservation

The clustering stage is similar to the deep embedded clustering [1], but we adopt cross entropy instead of K-L divergence to accelerate convergence speed. We apply cluster process on the concatenate layer of latent RNA representation and protein input data.

Our clustering algorithm is defined as cross entropy between distribution *P* and *Q*, where *Q* is the distribution of soft labels measured by Student’s *t* -distribution and *P* is the derivation of the target distribution from *Q*. Formally, the clustering loss is

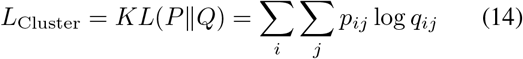

where *q*_*ij*_ is the soft label of embedded point *z*, which is defined as the similarity between *z*_*i*_ and cluster centre *µ*_*i*_ measured by Student’s *t* -distribution

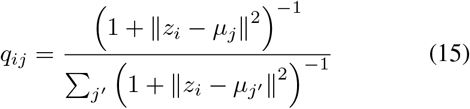

Meanwhile, *p*_*ij*_ is the target distribution computed by first raising *q*_*ij*_ to the second power and then normalizing by frequency per cluster:

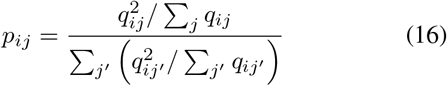

We adopt cross entropy in this stage instead of K-L divergence so that there are no division operations when calculating thus get a faster convergence during training. This training strategy can be seen as self-training, because the target distribution *P* is defined based on *Q*. We pretrain the stacked denoising ZINB model-based autoencoder before the concatenate stage and pre-train the CCA loss added model before clustering stage. The initialization of cluster centres is obtained by standard *k* -means clustering in the embedded feature space after pre-training of the denoising NB model-based autoencoder. As a result, the model has three components: the denoising NB model-based autoencoder, a concatenate layer and the clustering part. Then the objective function of scCTClust is

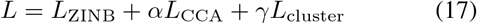

where *L*_ZINB_, *L*_CCA_ and *L*_Cluster_ are the ZINB loss functions of denoising the NB model-based autoencoder, CCA_loss and the clustering loss, respectively, and *α, γ >* 0 is the coefficient that controls the relative weights of the three losses. According to [7], this form of loss has the advantage that it can preserve the local structure of the data-generating distribution.

## 4. Conclusion

Nowadays, as multi-omics sequencing methods gradually became mature, more and more researchers focus on the downstream analysis over multi-omics sequencing data. Casting our sight on clustering analysis over CITE-seq data, we are facing not only problems RNA-seq data brings in but also how to integrate different modalities of data. To characterize the RNA-seq data obtained from droplet experiments, we used a ZINB based denoising autoencoder model. We introduced a cca_loss which is rooted in CCA to maximize the correlation between protein count matrix and latent representation of RNA data in order to correct clustering with protein information as auxiliary. Finally we conducted clustering process in a way similar to T-SNE but replace the dec_loss with cdec_loss defined in section.3 to accelerate convergence and improve clustering performance.

As the size of single cell sequencing datasets inflated at an amazing speed over the past few years, the import of more powerful dimension reduction methods became necessary in the domain of single cell clustering analysis. And with the gush of different type of single cell multiomics sequencing techniques, assorted multi-modal analyzing tools are needed eagerly. As a deep learning multimodal clustering method with effective dimension reduction function, it is not not surprising that scCTClust gained outstanding performance among state-of-art methods over the extensive CITE-seq datasets. But nobody is perfect, scCTClust occupied large memory space and is less stable than those methods without cca_loss as SVD and matrix inversion are needed during the optimization of cca_loss. And that unstableness is exactly what we endeavor to avoid to improve our method at present.

## Acknowledgments

We acknowledge anonymous reviewers for the valuable comments on the original manuscript.

